# Studies on Biochemical Changes and Prolactin Level Evaluation in Patients with Chronic Kidney Disease

**DOI:** 10.1101/677708

**Authors:** Foyzur Rahman, Faisal Kabir, Sawgat Rezwan

## Abstract

**Background:** Chronic kidney disease (CKD) is progressive loss in kidney function over a period of months or years. CKD is an internationally recognized public health problem affecting 5–10% of the world population and day-by-day the number of cases are increasing at an alarming rate. In CKD, raised levels of prolactin in blood may cause vascular derangements which might lead to worse cardiovascular consequences in CKD patients.

**Objectives:** To assess serum creatinine, hemoglobin (Hb), urea, red blood cell (RBC), protein creatinine ratio (PCR) and prolactin in CKD patients.

**Material and Methods:** This study included 110 patients, 61 were males and 49 were females and their age range 1 to more than 60 years. The control group also consisted of same number of participants as the patients; who were free from signs and symptoms of kidney disease and prolactin hormone disorders.

**Results:** The study shows that all the biochemical parameters in CKD patients were found to be significantly high compared with control group (P<0.001). Serum prolactin concentrations in CKD patients were also increased significantly compared with control group (P≤ 0.05). It was found that level of prolactin hormone secretion was higher in male CKD patients than male control.

**Conclusion:** Although males are more prone to chronic kidney disease, but the percentage of females is not negligible. All the biochemical parameters and prolactin level changed significantly in the CKD patients. It is interesting that in case of CKD, male’s prolactin secretion becomes higher.

## Introduction

Chronic kidney disease (CKD) is progressive damage in kidney function over a period of months or years ^[1]^. The symptoms of worsening kidney function are not specific, and might include feeling generally unwell and experiencing a reduced appetite ^[2]^. CKD is a long-term form of kidney disease; thus, it is differentiated from acute kidney disease (acute kidney injury) in that the reduction in kidney function must be present for over 3 months. CKD is an internationally recognized public health problem affecting 5–10% of the world population ^[3]- [4]^. The most common recognized causes of CKD are diabetes mellitus and high blood pressure. Other causes of CKD include idiopathic (i.e. unknown cause, often associated with small kidneys on renal ultrasound) and glomerulonephritis ^[5]-[6]^.

Previous professional guidelines classified the severity of CKD in five stages, with stage 1 being the mildest and usually causing few symptoms and stage 5 being a severe illness with poor life expectancy if untreated. But recent international guidelines reclassified CKD based on cause, glomerular filtration rate category (G1, G2, G3a, G3b, G4 and G5), and albuminuria category ^[7]^. Among these, glomerular filtration rate category is widely used.

CKD can be detected via simple biochemical tests ^[8]^. These tests include a creatinine-based estimate of the glomerular filtration rate (GFR), serum creatinine ^[2]^, RBC, urea etc. ^[9]^.

CKD is characterized by elevation of serum prolactin levels. Prolactin is a hormone secreted mainly by anterior pituitary gland. Main action of prolactin is to control breast development and lactation in women. The function of prolactin in men remains to be studied. Prolactin clearance is reduced in CKD, and its production is altered ^[10]^. In male CKD patients, hyperprolactinemia is associated with gynecomastia and sexual dysfunction ^[11]^.

The objective of this study was to find out the biochemical changes in patients with chronic kidney disease and compare the obtained results with the results of healthy individuals as control groups and also to assess the serum prolactin levels in CKD patients.

## Material and Methods

### 2.1 Case preparation

This is a cross sectional study done at Khwaja Yunus Ali Medical College Hospital in Enayetpur, Sirajganj. A number of 110 CKD patients (61 males and 49 females) aged 1 to more than 60 years were subjected to investigation. 110 normal individuals whose renal parameters were within normal limits and with no history of renal impairment in the past were also selected between the comparable age group who functioned as the control group (matched for age and sex). The sample population visited at the hospital from September, 2018 to November, 2018. The biochemical parameters serum creatinine, urea and protein creatinine ratio (PCR) were measured by automated chemistry analyzer Beckman coulter AU-480. Hemoglobin (Hb) and red blood cell (RBC) were measured by Sysmex XN-1000 analyzer. Automated Cobas e 411 Immunoassay analyzer was used to assess prolactin level. Approval from the institutional authority was taken prior to the study.

### 2.2 Statistical analysis

The data obtained from this study were analyzed with SPSS 21.0 program. Results were expressed as mean ± standard deviation (SD). Statistical significance was assessed by Independent-Samples T Test. The mean and standard deviation were calculated for all variables. These variables were compared between patients and healthy persons. The Independent-Samples T Test was used to compare these variables between patients and healthy persons. A P-value ≤ 0.05 was considered statistically significant.

## Results

At the time of the study, the majority of the patients (N=110) were adults (96.3%) and of male gender (55.45%), whereas the total number of female patients were 49 which constitutes 44.54% of the whole figure (Fig. 1).

**Fig. 1:**
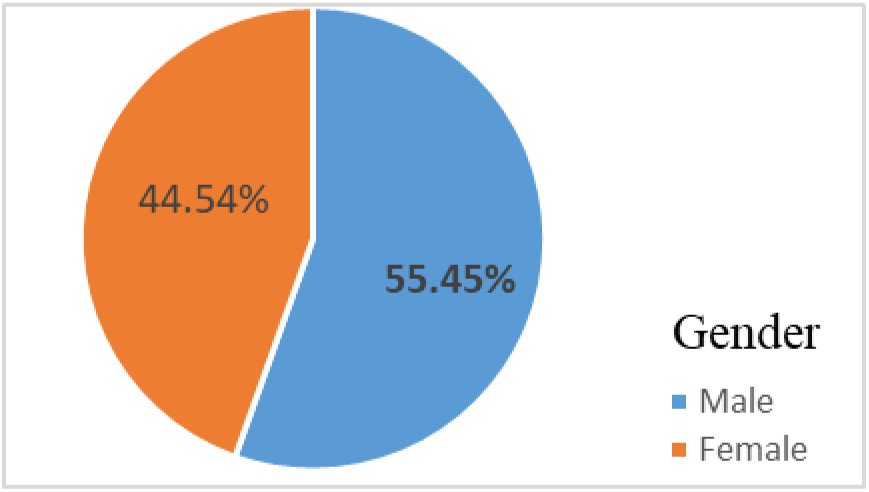
Pie chart showing gender distribution of CKD patients.

All the CKD patients were grouped into four (<20, 21-40, 41-60 and >60) based on age and it was found that most of the patients (42.72%) were aged from 41 to 60. Only 5 cases were below age 20 (Fig. 2).

**Fig. 2:**
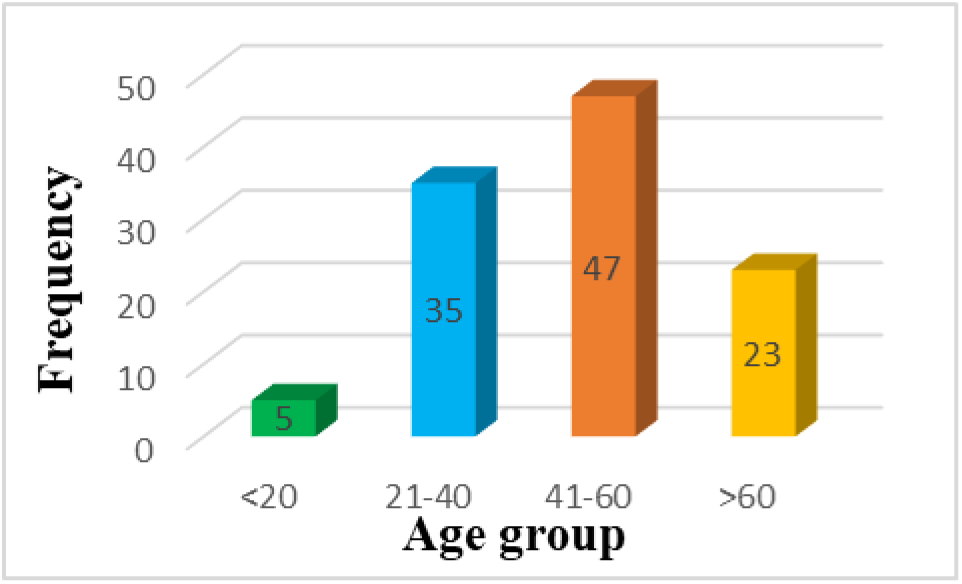
Number of patients of different age groups.

We estimated the glomerular filtration rate (eGFR) of all the CKD patients to determine how the renal function of the patients still have. It was found that, patients of every CKD stages were present in the study among them most of the patients (n=53) were of G5 stage. In total, 10 out of 110 came of the first three CKD stages (Table I).

**Table I:** eGFR level of cases in different stages of CKD.

| CKD Stages | Frequency |
| --- | --- |
| G1 | 2 |
| G2 | 1 |
| G3a | 7 |
| G3b | 16 |
| G4 | 31 |
| G5 | 53 |

The results in Table II demonstrated the level of biochemical parameters in both male and female in case of chronic kidney failure patients and control groups.

**Table II:** Comparison of biochemical parameters between CKD cases and controls (mean±SD).

| Biochemical Parameters | CKD cases (mean±SD) | Controls (mean±SD) | P value |
| --- | --- | --- | --- |
| Serum creatinine (umol/l) | 556.80±539.89 | 67.38±25.33 | <0.001** |
| Hb (mg/dl) | 9.24±2.14 | 15.22±1.82 | <0.001** |
| RBC (10<sup>12</sup>/L) | 3.35±0.77 | 4.00±0.69 | <0.001** |
| Urea (mmol/l) | 24.15±14.13 | 5.00±1.85 | <0.001** |
| PCR (mg/mmol) | 1012.90±907.23 | 35.46±10.26 | <0.001** |

\*\*significant.

Among the parameters, level of serum creatinine, blood urea and PCR were found to be significantly increased in mean values in CKD cases as compared to the controls with a P value of <0.001, whereas hemoglobin and RBC were significantly decreased.

From the above Table III we can demonstrate that, the mean average level of renal parameters is statistically less significant i.e. no difference according to age i.e. p-value was > 0.05. Mean urea, creatinine and PCR are comparatively higher in younger age group but differences are not significant.

**Table III:** Age differentials in parameters of CKD patients.

| Age group→ | <20 yrs.<br>(n=5) |  | 21-40 yrs.<br>(n=35) |  | 41-60 yrs.<br>(n=47) |  | >60 yrs.<br>(n=23) |  | P value |
| --- | --- | --- | --- | --- | --- | --- | --- | --- | --- |
| Parameters<br>↓ | Mean | SD | Mean | SD | Mean | SD | Mean | SD |  |
| Creatinine | 344.59 | 305.19 | 754.94 | 700.37 | 468.14 | 482.53 | 482.60 | 295.05 | 0.15 |
| Urea | 15.05 | 8.28 | 26.47 | 17.64 | 20.98 | 11.20 | 29.07 | 12.65 | 0.27 |
| Hb | 10.22 | 0.71 | 8.72 | 2.55 | 9.46 | 2.13 | 9.36 | 1.51 | 0.08 |
| RBC | 3.68 | 0.43 | 3.14 | 0.95 | 3.51 | 0.70 | 3.31 | 0.56 | 0.07 |
| PCR | 659.46 | 161.64 | 1035.20 | 794.08 | 1129.04 | 1157.27 | 818.45 | 448.13 | 0.47 |

None of the biochemical parameters showed significant sex differentials (Table IV). Except Hb, level of serum creatinine, RBC, urea and PCR for male are higher than female but the difference is not significant.

**Table IV:** Sex differentials in the parameters of CKD patients.

| Parameters | Male (n=61) |  | Female (n=49) |  | P value |
| --- | --- | --- | --- | --- | --- |
|  | Mean | SD | Mean | SD |  |
| Creatinine | 749.29 | 607.74 | 317.18 | 307.45 | 0.98 |
| RBC | 3.29 | 3.29 | 3.44 | 0.73 | 0.85 |
| Hb | 9.35 | 2.28 | 9.10 | 1.96 | 0.09 |
| Urea | 28.11 | 14.66 | 19.22 | 11.85 | 0.67 |
| PCR | 1255.47 | 1088.93 | 710.92 | 467.93 | 0.77 |

### Serum Prolactin

Serum prolactin level was measured for the 20 patients with CKD using fully automated Cobas e 411 Immunoassay analyzer. Among the 20 CKD patients, 15 patients had raised serum prolactin levels (Table V).

**Table V:**
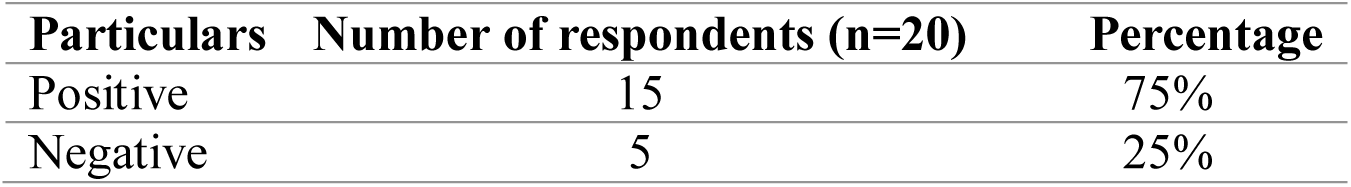
Serum prolactin percentage

Patients with kidney disease secret much prolactin hormone than the healthy individuals and it was found that the mean was about more than two times higher in CKD patients (Fig. 3).

**Fig. 3:**
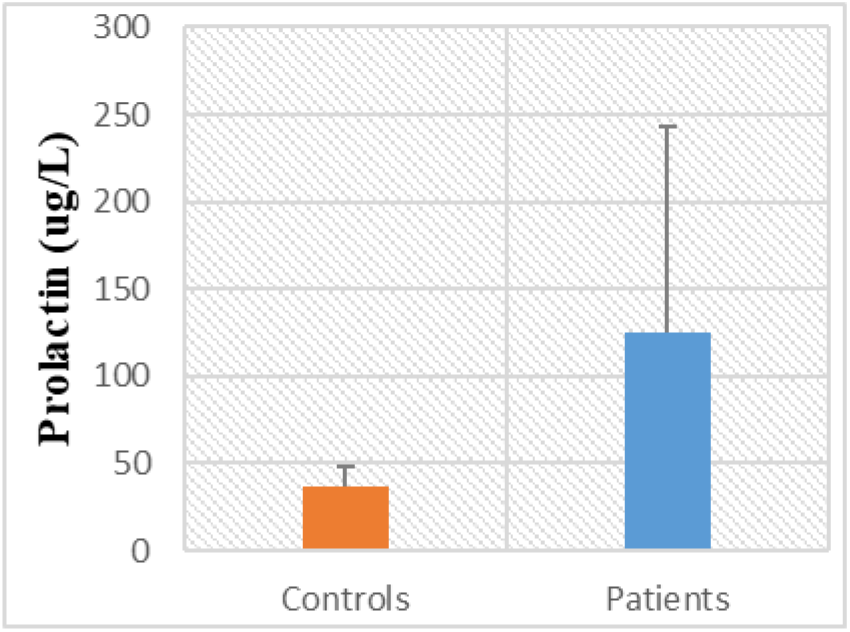
The statistical histogram of prolactin for all participants (patients and controls) with standard deviation.

We also compared prolactin secretion between male CKD patients and male controls and found a significant increase of prolactin secretion in CKD patients (Fig. 4).

**Fig. 4:**
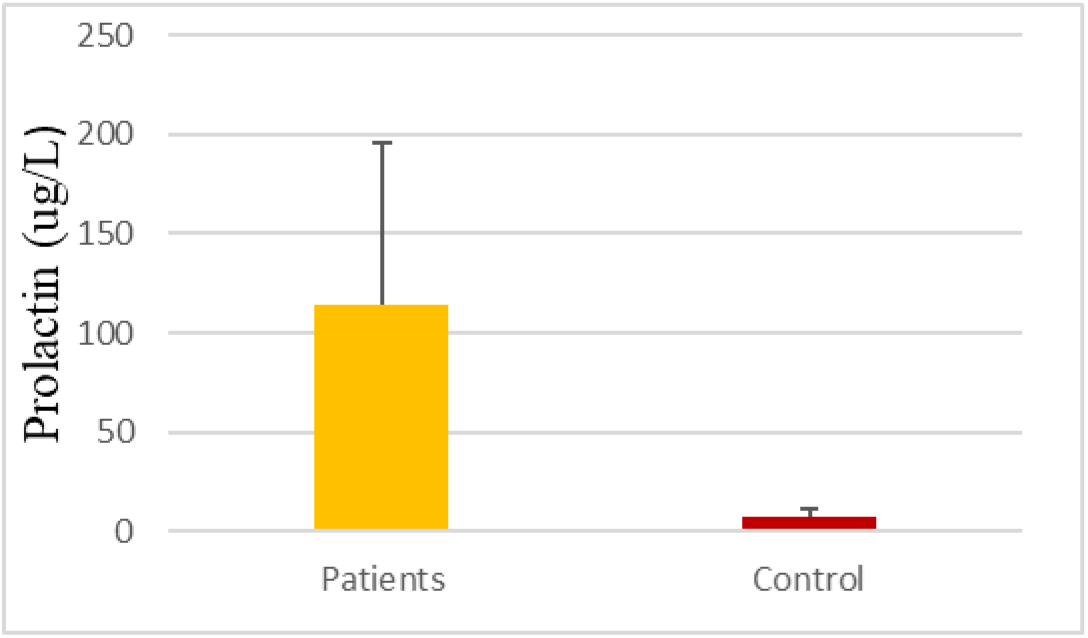
The statistical histogram of prolactin for CKD male patients and male control with standard deviation.

According to statistical analysis of data using t-test, there is a significant association between increased serum prolactin levels and presence of CKD (Table VI).

**Table VI:** t-test, serum prolactin levels (ug/L) significance of association between chronic kidney disease and increased serum prolactin levels.

| Hyperprolactinemia | Mean | Standard deviation | Statistical inference |
| --- | --- | --- | --- |
| Positive (n=15) | 160.88 | 112.44 | t= -5.13 |
| Negative (n=5) | 11.64 | 2.85 | df= 14.05 |
|  |  |  | 0.000<0.05 |
|  |  |  | Significant |

## Discussion

The present work was conducted in order to identify biochemical changes and prolactin secretion in CKD patients in comparison with controls, with emphasis on differences which may have had an impact on diagnostic or therapeutic approaches. Several studies correspond that chronic kidney disease typically increases with age ^[8], [12]^ which supports this study. Older adults are, therefore, at increased risk for chronic kidney disease ^[13]^. It is found that the female gender is associated with slower progression of chronic kidney disease. It also supports the present study in which 44.54% of the total patients are female (Fig. 1).

Among the total patients, 48.18% patients came to the hospital with severe illness with poor life expectancy if untreated (CKD stage G5). In addition, another 31 cases were of G4 stage (Table I). These results mean that people of our country are not aware of kidney disease or they do not care about their renal health.

Among the parameters, the mean average level of serum creatinine, blood urea and PCR in patients were measured 556.80±539.89 umol/l, 24.15±14.13 mmol/l and 1012.90±907.23 mg/mmol respectively, which were significantly higher than the controls (67.38±25.33 umol/l, 5.00±1.85 mmol/l and 35.46±10.26 mg/mmol respectively). Although changed significantly, the mean average level of hemoglobin and RBC in patients (9.24±2.14 mg/dl and 3.35±0.77×10^12/L respectively) were decreased than the controls (15.22±1.82 mg/dl and 4.00±0.69×10^12/L respectively). These changes are supported by several studies-Bhan et.al. ^[14]^, Shah et.al. ^[15]^. According to different age groups, mean urea, creatinine and PCR are comparatively higher in younger age and male group but differences are not significant.

In the study, blood prolactin level was measured in 20 patients. Among them, 75% patients showed an increase in prolactin level known as Hyperprolactinemia (Table V). Hyperprolactinemia is defined as a serum prolactin level above the normal range (25ng/mL in women and 15ng/mL in men) ^[1]^. The secretion of prolactin is influenced by various substances including monoamines, endogenous opiates and can be altered in uremia ^[16]^.

The mean prolactin level in patients is significantly higher than the controls (Fig. 3). This may be due to the reduced renal clearance or increased production of prolactin ^[17]^. We also found a significant increase of prolactin secretion in CKD male patients comparing to the male controls (Fig. 4). This may be due to increased release of prolactin from the anterior pituitary, due to increased production of prolactin in immune system cells or due to the reduced renal clearance ^[17]^, ^[18]^. In addition, our results showed that the mean of prolactin in the male is less than the female mean.

## Conclusion

Although males are more prone to chronic kidney disease, but the percentage of females is not negligible. All the biochemical parameters and prolactin level changed significantly in the CKD patients. It is interesting that in case of CKD, prolactin secretion in male patients becomes higher.

## Acknowledgement

We are grateful to the chairman of the Trustee Board and Director of Khwaja Yunus Ali Medical College Hospital to give permission for conduction of this study.

